# Vaginal microbiome-host interactions modeled in a human vagina-on-a-chip

**DOI:** 10.1101/2022.03.20.485048

**Authors:** Gautam Mahajan, Erin Doherty, Tania To, Arlene Sutherland, Jennifer Grant, Abidemi Junaid, Aakanksha Gulati, Nina Teresa LoGrande, Zohreh Izadifar, Sanjay Sharma Timilsina, Viktor Horváth, Roberto Plebani, Michael France, Indriati Hood-Pishchany, Seth Rakoff-Nahoum, Douglas S. Kwon, Girija Goyal, Rachelle Prantil-Baun, Jacques Ravel, Donald E. Ingber

**Affiliations:** Wyss Institute for Biologically Inspired Engineering at Harvard University, Boston, MA 02115, USA; Institute for Genome Sciences and Department of Microbiology & Immunology, University of Maryland School of Medicine, MD, Baltimore, USA; Division of Infectious Diseases and Division of Gastroenterology, Department of Pediatrics, Boston Children’s Hospital and Harvard Medical School, 300 Longwood Avenue, Boston, 02115, Massachusetts, USA; Ragon Institute of MGH, MIT, and Harvard, Massachusetts General Hospital, Harvard Medical School, Boston, MA, USA; Division of Infectious Diseases, Massachusetts General Hospital, Boston, MA, USA; Vascular Biology Program and Department of Surgery, Boston Children’s Hospital and Harvard Medical School, Boston, MA 02115, USA; Harvard John A. Paulson School of Engineering and Applied Sciences, Harvard University, Cambridge, MA 02139, USA

## Abstract

**Background:** A dominance of non-iners *Lactobacillus* species in the vaginal microbiome is optimal and strongly associated with gynecological and obstetric health, while the presence of diverse obligate or facultative anaerobic bacteria and a paucity in *Lactobacillus* species, similar to communities found in bacterial vaginosis (BV), is considered non-optimal and associated with adverse health outcomes. Various therapeutic strategies are being explored to modulate the composition of the vaginal microbiome; however, there is no human model that faithfully reproduces the vaginal epithelial microenvironment for preclinical validation of potential therapeutics or testing hypotheses about vaginal epithelium-microbiome interactions.

**Results:** Here, we describe an organ-on-a-chip (Organ Chip) microfluidic culture model of the human vaginal mucosa (Vagina Chip) that is lined by hormone-sensitive, primary vaginal epithelium interfaced with underlying stromal fibroblasts, which sustains a low physiological oxygen concentration in the epithelial lumen. We show that the Vagina Chip can be used to assess colonization by optimal *L. crispatus* consortia as well as non-optimal *Gardnerella vaginalis*-containing consortia, and to measure associated host innate immune responses. Co-culture of the *L. crispatus* consortia was accompanied by maintenance of epithelial cell viability, accumulation of D- and L-lactic acid, maintenance of a physiologically relevant low pH, and down regulation of proinflammatory cytokines. In contrast, co-culture of *G. vaginalis-*containing consortia in the Vagina Chip resulted in epithelial cell injury, a rise in pH, and upregulation of proinflammatory cytokines.

**Conclusion:** This study demonstrates the potential of applying human Organ Chip technology to create a preclinical model of the human vaginal mucosa that can be used to better understand interactions between the vaginal microbiome and host tissues, as well as to evaluate the safety and efficacy of live biotherapeutics products.

## INTRODUCTION

There is growing recognition of the pivotal role the microbiome plays in regulation of vaginal health and disease[1]. Vaginal microbiota dominated by *Lactobacillus* species such as *L. crispatus, L. gasseri,* and *L, jensenii* are considered to be a hallmark of an optimal microbiome found within the female reproductive tract and are associated with positive health outcomes[2]. These lactobacilli modulate the vaginal microenvironment through their metabolic actions and by production of bioactive compounds (e.g., D- and L-lactate[3, 4] and bacteriocins[5, 6]), which collectively contribute to protect the vagina against pathogenic bacteria. In particular, lactobacilli produce copious amount of lactic acid acidifying the vagina to pH < 4.5. At concentration found in the vaginal microenvironment, lactic acid has been shown to have antimicrobial, antiviral and anti-inflammatory properties[4]. In contrast, non-optimal vaginal microbiota are characterized by a paucity of lactobacilli and the presence of a wide array of strict and facultative anaerobes, often including *Gardnerella vaginalis*[7–9]. These non-optimal vaginal microbiota composed of diverse anaerobe-dominant consortia are reminiscent of those associated with the conditions bacterial vaginosis (BV)[9] and have been associated with increased susceptibility to and transmission of sexually transmitted infections[10], as well as increased risk of pelvic inflammatory disease[11], maternal infections[12], and preterm birth[7, 13] which is the second major cause of neonatal death across the world[14].

Given the key role that the microbiome appears to play in regulating vaginal health and disease, there is renewed interest in exploring the use of live biotherapeutic products to modulate the composition and function of the vaginal microbiome and thereby treat or prevent BV and its associated sequelae[15, 16]. Recently, promising results were obtained in a phase 2b clinical trial in which a live biotherapeutic product containing a single *L. crispatus* strain (LACTIN-V) was shown to decrease risk of recurrent BV when administered after standard of care metronidazole treatment[15]. However, the development of new therapeutic strategies to treat diseases and disorders of the female reproductive tract has been hampered by the lack of relevant human vaginal epithelium models. This is a critical need as animal models are of limited use in research done to study host-microbiota interactions in the vaginal space, because of the major physiological, anatomical and microbial differences present in these models compared compared to the human vagina[17].

Most of our knowledge of the composition and function of the vaginal microbiome comes from genomic and metagenomic analysis of clinical samples. However, it is difficult to study how the vaginal microbiome interactions with human vaginal epithelium under controlled conditions in a physiologically relevant microenvironment. Bacteria can be co-cultured with cultured human cells or more complex *in vitro* models, such as organoids and transwells, but they only support short-term (< 1 day) co-culture with living microbes before they lead to bacteria overgrowth and cell death[18–21]. More importantly, these static *in vitro* models fail to recapitulate physiologically relevant tissue-tissue interfaces and other microenviromental cues (e.g, epithelial-stromal interactions, air-liquid interface, dynamic fluid flow, etc.) that are critical for recapitulation of organ-level physiology and pathophysiology[21]. A similar challenge has been successfully overcome in context of the human gut microbiome using organ-on-a-chip (Organ Chip) microfluidic culture technology [22, 23], which has been shown to enable sustained culture of complex living microbiota in contact with human intestinal epithelium for at least 5 days *in vitro*[19]. Thus, in the present study, we set out to leverage Organ Chip technology to create a microfluidic culture device lined by human vaginal epithelium interfaced with stromal fibroblasts, and to explore whether it can be used to study host tissue interactions microbial consortia dominated by *L. crispatus* versus *G. vaginalis.* Here, we show that *L. crispatus* consortia stably engraft in the Vagina Chip, establish an acid pH, produce both D- and L-lactate, and down-regulate proinflammatory cytokines. Moreover, culture of *G. vaginalis-containing* microbial consortia or *G. vaginalis* alone on-chip increased pH and secretion of inflammatory cytokines, and resulted in epithelial cell injury. Thus, the Vagina Chip may represent a human *in vitro* preclinical model that can be used to advance host-microbiome research and accelerate development of microbiome-targeted therapeutics including live biotherapeutic products.

## METHODS

### Human Vagina Chip Culture

Microfluidic two-channel co-culture Organ Chip devices were obtained from Emulate Inc. (Boston, MA). The apical channel (1 mm wide × 1 mm high) and basal channel (1 mm wide × 0.2 mm high) are separated by the porous membrane (7 μm diameter pores) along their length (16.7 mm). For activation, both channels were filled with 0.5 mg/mL ER1 solution in ER2 buffer (Emulate Inc.) and placed under UV light for 20 minutes followed by washing with ER2 buffer and phosphate-buffered saline (PBS). Before cell seeding, the apical channel was incubated with collagen IV (30 μg/mL) from human placenta (Sigma, cat. no. C7521) and collagen I (200 μg/mL) from rat tail (Corning, cat. no. 354236) in DMEM (ThermoFisher, cat. no. 12320-032) at 37°C with 5% CO_2_ for 2-3 hours. The basal channel was incubated with collagen I (200 μg/mL) from rat tail (Corning, cat. No. 354236) and poly-L-lysine (15 μg/mL) (ScienCell Research Laboratories, cat. no. 2301) in DMEM (ThermoFisher, cat. no. 12320-032) at 37°C with 5% CO_2_ for 2-3 hours.

Primary human vaginal epithelial cells (Lifeline Cell Technology, cat. no. FC-0083; donors 05328 and 04033) were expanded in 75-cm^2^ tissue-culture flasks using vaginal epithelium growth medium (Lifeline Cell Technology, cat. no. LL-0068) to 60-70% confluency. Primary human uterine fibroblasts (ScienCell Research Laboratories, cat. no. 7040) were expanded in 75-cm^2^ tissue-culture flasks coated with poly-L-lysine (15 μg/mL, ScienCell Research Laboratories, cat. no. 2301) using fibroblast growth medium (ScienCell Research Laboratories, cat. no. 7040) to 60-70% confluency.

To create the human Vagina Chip, fibroblasts (1 × 10^6^ cells/mL) were seeded first in the basal channel by inverting the chip for 1 hour in human fibroblast growth medium. Chips were inverted again, and human vaginal epithelial cells (3 × 10^6^ cells/mL) were seeded in the apical channel for 4 hours in human vaginal growth medium. The chips were incubated at 37°C with 5% CO_2_ overnight under static conditions. The basal channel was continuously perfused with fibroblast growth medium using the Zoe culture module (Emulate) at a volumetric flow rate of 40 μL/h. The apical channel was intermittently perfused with vaginal epithelium growth medium by changing the flow rate in the apical channel from 0 to 40 μL/h for 4 hours per day by using the Zoe culture module. After 5-6 days, the apical medium was replaced with Hank’s Balanced Salt Solution (HBSS; ThermoFisher, cat. no. 14025092) and the basal medium was replaced with in-house differentiation medium (see below) for 8 days following same intermittent and continuous perfusion regime, respectively. The HBSS was further replaced with customized HBSS Low Buffer/+Glucose (HBSS (LB/+G)) for 2 days followed by 3 days with microbial co-culture as described below.

Customized HBSS (LB/+G) medium is composed of 1.26 mM calcium chloride (Sigma, cat. no. 499609), 0.49 mM magnesium chloride hexahydrate (Sigma, cat. no. M2393), 0.41 mM magnesium sulfate heptahydrate (Sigma, cat. no. M2773), 5.33 mM potassium chloride (Sigma, cat. no. P5405), 0.44 mM potassium phosphate monobasic (Sigma, cat. no. P5655), 137.93 mM sodium chloride (Sigma, cat. no. S5886) and 5.56 mM D-glucose (Sigma, cat. no. G7021).

In-house differentiation medium is composed of DMEM (ThermoFisher, cat. no. 12320-032), Ham’s F12 (ThermoFisher, cat. no. 11765-054), 4 mM L-glutamine (ThermoFisher, cat. no. 25030081), 1 μM hydrocortisone (ThermoFisher, cat. no. H0396), 1X Insulin-Transferrin-Ethanolamine-Selenium (ITES; Lonza, cat. no. 17-839Z), 20 nM triiodothyronine (Sigma, cat. no. T6397), 100 μM O-phosphorylethanolamine (Sigma, cat. no. P0503), 180 μM adenine (Sigma, cat. no. T6397), 3.2 mM calcium chloride (Sigma, cat. no. 499609), 2% heat inactivated fetal bovine serum (FBS; ThermoFisher, cat. no. A3840001), 1% penicillin -streptomycin (ThermoFisher, cat. no. 15070063) and 4 nM β-Estradiol (Sigma, cat. no. E2257).

### Immunofluorescence Microscopy

The Vagina Chips were fixed with 4% paraformaldehyde (Alfa Aesar, stock no. J61899) for 30 minutes at room temperature and washed with phosphate buffered saline (PBS). The channels were filled with 2% agarose (Lonza, cat. no. 50302) and the whole chip was embedded in O.C.T. compound (Fisher Scientific, cat. no. 23-730-571) and stored at −80°C until sectioning. Chips were cryosectioned at a thickness of 50 μm on a cryostat (Leica CM3050 S). The cryosections were then permeabilized using 0.1% Triton-X (Sigma-Aldrich, SKU no. X100) in PBS, blocked with 5% goat serum (Life Technologies, cat no. 16210072) in 0.01% Triton-X in PBS for 1 hour at room temperature, and then incubated at 4°C overnight with primary antibodies against CK13 (Abcam, cat. no. ab92551 at 1:200 dilution), CK14 (Abcam, cat. no. ab119695 at 1:200 dilution), E-cadherin (Abcam, cat. no. ab40772 at 1:200 dilution), ZO-1 (Abcam, cat. no. ab276131 at 1:40 dilution), Involucrin (Abcam, cat. no. ab68 at 1:200 dilution), DSG1 (Abcam, cat. no. ab12077 at 1:400 dilution), and DSG3 (Abcam, cat. no. ab231309 at 1:400 dilution). The sections were washed 3 times with PBS, and then incubated with secondary antibody (Abcam, cat. no. ab150077) at a dilution of 1:500 for 1 hour at room temperature. Some sections also were incubated with directly labeled fluorescent with Alexa Fluor® antibodies against CK5 (Abcam, cat. no. ab193894) or CK15 (Abcam, cat. no. ab194065) or phalloidin (Invitrogen, cat. no. A22287) in the dark at 4°C. Vagina Chip sections were stained with as-received Eosin Y solution (Abcam, cat. no. ab246824), which fluoresces under blue-green excitation, for 30 seconds at room temperature to obtain pseudo-H&E staining. All stained sections were counterstained with 4’,6-diamidino-2-phenylindole (DAPI; Invitrogen, cat. no. D1306) at a concentration of 1 μg/mL for 15 minutes at room temperature and mounted using ProLong Glass Antifade Mountant (ThermoFisher, cat. no. P36980). Images were acquired with an inverted laser-scanning confocal microscope (Leica SP5 X MP DMI-6000) and processed using ImageJ/Fiji. Pseudo H&E images were processed using ImageJ/Fiji and MATLAB (Mathworks) using a previously published method[24].

### Barrier Permeability

Cascade blue (Invitrogen, cat. no. OC3239) was added to apical channel media at a concentration of 50 μg/mL. Effluent from the apical and basal channels were collected and measured for fluorescence intensity at an excitation wavelength of 380 nm and an emission wavelength of 420 nm using a multi-mode plate reader (BioTek NEO). The apparent permeability (P_app_) was calculated using the equation as previously reported[25]: P_app_ = (V_r_ * C_r_)/A * t *(C_d-out_ * V_d_ + C_r_ * V_r_) / V_d_ + V_r_)), where V_r_ is the volume of the receiving channel, V_d_ is the volume of the dosing channel, A is the area of the co-culture membrane, t is the total time of effluent flow, C_r_ is the measured concentration of Cascade Blue in the receiving channel effluent, and C_d-out_ is the measured concentration of Cascade Blue in the dosing channel effluent.

### RT-qPCR

Total RNA was extracted from vaginal epithelial cells from pre-differentiated (day 5 of expansion) and differentiated (day 10 of differentiation; exposed to 0.4 nM and 4 nM of β-estradiol for 10 days) Vagina Chips using QIAzol lysis reagent (Qiagen, cat. no. 79306). Complimentary DNA was synthesized using a SuperScript VILO MasterMix (Invitrogen, cat. no. 11755-500). The cellular gene-expression levels were determined using RT–qPCR, according to the TaqMan fast advanced master mix (ThermoFisher, cat. no. 4444964) with 20 μL of a reaction mixture containing gene-specific primers (ThermoFisher) for estrogen receptor (ESR1, Hs01046816), progesterone receptor (PGR, Hs01556702), phosphoenolpyruvate carboxykinase 1 (PCK1, Hs00159918), claudin 17 (CLDN17, Hs01043467), glucagon receptor (GCGR, Hs00164710), keratin15 (KRT15, Hs00951967) and zonula occludens-1 (ZO-1, Hs01551871). The expression levels of the target genes were normalized to GAPDH (Hs04420632).

### Computational Simulations

Using COMSOL Multiphysics 5.5 (COMSOL, Inc.) a two-dimensional model of two-channel microfluidic device was developed. The co-culture window was used to model the oxygen gradient with 80 μm epithelium and 50 μm stroma in the apical and basal channel, respectively. The apical and basal PDMS blocks are 3.5 mm and 1 mm thick respectively, and the PDMS membrane is 50 μm thick. The 2D oxygen distribution was simulated by coupling laminar flow with dilute species transport. The oxygen-saturated medium was fed through the inlet at the flow rate of 40 μl/hr and goes out of the outlet after being partially consumed by the cells via aerobic respiration. Oxygen consumption by the epithelium and stroma was simulated using Michaelis-Menten-type kinetics. Navier-Stokes equations for incompressible flow were used to simulate fluid flow, and Fick’s second law was used to simulate oxygen transport through the PDMS, culture medium, epithelium, and stroma. Steady-state and time-dependent simulations were performed at 37 °C and with 145 mm Hg atmospheric pO2 to simulate the conditions in the cell culture incubator. The entire vagina chip contained 145 mm Hg atmospheric pO2 at t=0 min and the time-dependent model was simulated for 200 min of continuous flow.

### Isolation and Selection of *L. crispatus* Strains (C0006A1, OC1, OC2 and OC3)

As recently reported, vaginal microbiota dominated by *Lactobacillus* spp. comprise of multiple strains of the same species[26]. Consequently to mimic the ecology of these optimal vaginal microbiota, three *L. crispatus* multi-strain consortia were reconstructed that contain *L. crispatus* isolates cultivated from women with stable *L. crispatus* dominated microbiota who participated in the UMB-HMP study[27]. One optimal consortium (OC1) contained four *L. crispatus* strains (C0175A1, C0124A1, C0112A1 and C0059A1), while two of the optimal consortia (OC2 and OC3) contained three *L. crispatus* strains (OC2: C0175A1, C0124A1 and C0059A1 and OC3: C0175A1, C0112A1 and C0006A1); C0006A1 contains a single strain that is also found within OC3 consortium.

### Isolation and Selection of *Gardnerella* Strains and Consortia (BVC1 and BVC2)

In non-optimal vaginal microbiota, *Gardnerella* species are typically found as dominant bacteria[7–9] accompanied by other frequent taxa such as *Prevotella* species and *Atopobium* species[2]. To mimic the ecology of non-optimal vaginal microbiota, two dysbiotic consortia (BVC1 and BVC2) were reconstructed from isolates cultivated from women with asymptomatic BV. The first contained complex consortia of taxa (BVC1: *G. vaginalis* E2, *G. vaginalis* E4, *P. bivia BHK8,* and *A. vaginae)* and second contained two strains of Gardnerella (BVC2: *G. vaginalis* E2 and E4). Recent studies have highlighted genomic diversity among Gardnerella spp. and the co-existence of multiple strains and species within an individual. The Gardnerella isolates used in this study were selected because they represent distinct genomic groups, exhibit phenotypic diversity in vitro, and were co-resident, meaning that they were co-isolated from a single participant in the UMB-HMP study[27]. P. bivia and A. vaginae are prevalent species in Lactobacillus-deficient vaginal microbiota. The two strains used in this study were co-resident, isolated from a single participant in the Females Rising Through Education Support and Health study[28].

### Construction of the Multi-strain *L. crispatus* Consortia and Inoculation in the Vagina Chip

Each unique *L. crispatus* strain was grown overnight at 37°C in De Man, Rogosa and Sharpe (MRS) broth (Fisher Scientific, cat. no. 288210) under complete anaerobic conditions (83% N_2_, 10% CO_2_, 7% H_2_) in an anaerobic chamber. Subcultures were made from overnight cultures and once mid-logarithmic phase was reached, aliquot stocks were made and frozen at −80°C with 16% sterile glycerol (MP Biomedicals, cat. no. 76019-966). To enumerate colony forming units (CFU)/mL in stocks, a single aliquot was thawed and spread plated on MRS agar (Hardy, cat. no. G117) under anaerobic conditions. Colonies were counted after 48 hours of incubation at 37°C and CFU/mL was calculated for stocks of each strain.

To generate consortia inoculum, required volumes from stocks of each strain were calculated in order to create equal *L. crispatus* strain cell density per 1 mL of inoculum. Cells were washed, spun, and resuspended in 1 mL of HBSS (LB/+G) and kept on ice. The apical channel of the Vagina Chip was inoculated with ~10^5^ CFU of prepared *L. crispatus* consortia on day 11 of differentiation and cultured for 72 hours. The chips were incubated statically at 37°C and 5% CO_2_ for first 20 hours of culture before starting the flow using the Zoe culture module. The basal channel was continuously perfused with in-house differentiation medium and apical channel was perfused for 4 hours per day with customized HBSS (LB/+G) medium at a volumetric flow rate of 40 μL/hr. Non-adherent bacterial CFU were quantified by measuring their presence in chip effluents collected at 24-, 48- and 72-hours post-inoculation and adherent bacteria were measured within epithelial tissue digests at 72 hours.

### Culture of a non-optimal *Gardnerella vaginalis* containing Consortium in the Vagina Chip

Two *G. vaginalis* strains (*G. vaginalis* E2 and *G. vaginalis* E4) and two other anaerobic bacteria found in non-optimal microbiota of patients with BV (*Prevotella bivia* BHK8, and *Atopobium vaginae*) were grown individually in Peptone, Yeast, and Tryptone (with hemin and vitamin K1) broth at 37°C under complete anaerobic conditions (83% N_2_, 10% CO_2_, 7% H_2_) in an anaerobic chamber. Subcultures were made from overnight cultures and once mid-logarithmic phase was reached, aliquot stocks were made and frozen at −80°C with 16% sterile glycerol (MP Biomedicals, cat. no. 76019-966). To enumerate CFU/mL in stocks, a single aliquot was thawed, serial diluted and spread plated on Brucella blood agar (with hemin and vitamin K1) (Hardy, cat. no. W23) under anaerobic conditions. Colonies were counted after 72 hours of incubation at 37°C and CFU/mL was calculated for stocks of each strain.

We then generated two consortia: one containing two *G. vaginalis* strains along with *P. bivia* BHK8 and *A. vaginae* species (BV Consortium 1, BVC1) and the other containing only *G. vaginalis* E2 and *G. vaginalis* E4 (BVC2). To generate these consortia, required volumes from stocks of each of the four bacterial strains were calculated in order to create equal strain cell density per 1 mL of inoculum. Cells were washed, spun, and resuspended in 1 mL of HBSS (LB/+G) and kept on ice. The apical channel was inoculated with ~10^6^ CFU of prepared BVC1 or BVC2 consortia and then chips were incubated statically at 37°C and 5% CO_2_ for 20 hours before starting the flow using the Zoe culture module. The basal channel was continuously perfused with in-house differentiation medium and apical channel was perfused for 4 hours per day with customized HBSS (LB/+G) medium at a volumetric flow rate of 40 μL/hr.

### Bacterial Enumeration from Vagina Chip Co-Culture

To enumerate all cultivable bacteria in the effluents, effluent samples (50 μL) were collected at 24, 48, and 72 hours, diluted with glycerol to a final concentration of 16%, and frozen at −80°C. *L. crispatus* samples were spread plated on MRS agar under complete anaerobic conditions. After 48 hours of incubation, colonies were counted, and CFU/mL was calculated for each sample. Effluent samples from the Vagina Chips containing BVC2 and BVC1 consortia were plated on Brucella blood agar (with hemin and vitamin K1) (Hardy, cat. no. W23) at 37°C under complete anaerobic conditions. After 72 hours of incubation, colonies were counted, and CFU/mL was calculated for each sample. To enumerate all cultivable bacteria engrafted in the Vagina Chip, the whole cell layer was digested for 3 hours with digestion solution containing 1 mg/mL of collagenase IV (Gibco, cat. no. 17104019) in TrypLE (ThermoFisher, cat. no. 12605010). Cell layer digest was then diluted with glycerol to a final concentration of 16% and frozen at −80°C. Digestion samples were processed in the same way as effluent samples and CFU/mL was calculated for each chip digest.

### Strain ratio analysis

DNA was extracted using the Qiagen AllPrep PowerViral DNA/RNA extraction kit (Qiagen; Hilden, Germany; Cat. 28000-50) from a 200 μL aliquot of the *L. crispatus* consortia inocula and from 200 μL of vaginal epithelial tissue digests after 72 hours of co-culture with each of the *L. crispatus* consortia. Four co-culture replicates were performed for each L. crispatus consortia. Following DNA extraction Illumina shotgun sequence libraries were prepared using the Kapa HyperPrep kit according to manufacturer specifications (Roche; Basen, Switzerland). Libraries were sequenced on an Illumina NovaSeq S4 flow cell (Illumina; San Diego, CA) yielding on average 45 million (range: 37.6-67.6 million) pairs of 150bp reads. Human reads were identified and removed using BMtagger[29]. Sequence datasets contained on average, 97.4% human reads (range: 96.3-98.3%). No human reads were identified in the inocula. Ribosomal RNA sequence reads were removed using sortmeRNA[30] (version 2.1) and the remaining reads were subjected to quality filtering and trimming usin fastp[31] (version: 0.21, sliding window size: 4bp, minimum average q-score: 20). The relative abundance of each *L. crispatus* strain in the samples was estimated by mapping the sequence reads to a database containing strain-specific marker genes. To build this database, single-copy genes uniquely present in the genomes of individual *L. crispatus* strains used in the experiments were identified using OrthoMCL[32] (all-versus-all BLAST e-value threshold 10^-5^, 70% percent identity, 70% overlap). Reads were mapped to the marker gene database using Bowtie2[33] and per gene coverage was estimated using SAMtools[34]. *L. crispatus* strain composition was determined using the median coverage of each strain’s marker genes relative to sum of median coverage values for all strains in the consortia. Sequence datasets were deposited in the Short Read Archive (##########). *L. crispatus* whole genome sequences have been deposited in genbank (#######). The *L. crispatus* strain specific marker gene database and all code used in the analysis of these data are available at: github.com/ravel-lab/#####).

### Lactate and pH

For lactate analysis, samples from apical effluent were collected at every 24-hour timepoint of the experiment and briefly equilibrated under anaerobic conditions (83% N_2_, 10% CO_2_, 7% H_2_) in an anaerobic chamber. Cells in each sample were pelleted and supernatant was collected then stored at 4°C. D-lactate and L-lactate concentrations were measured separately in each sample using BioAssay Systems EnzyChrom Lactate Assay Kits (cat. no. EDLC-100 and ECLC-100 respectively) according to the manufacturer’s protocol. During effluent collection, pH was measured using pH paper (Micro Essential, Hydrion 325) for all chips.

### Analysis of Cytokines and Chemokines

Samples (100 μL) of the apical effluents from Vagina Chips were collected and analyzed for a panel of cytokines and chemokines, including TNF-α, INF-y, IL-1α, IL-1β, IL-10, IL-8, IL-6, MIP-1α, MIP-1β, IP-10, TGF-β and RANTES using custom ProcartaPlex assay kits (ThermoFisher Scientific). The analyte concentrations were determined using a Luminex 100/200 Flexmap3D instrument coupled with the Luminex XPONENT software.

### Statistical analysis

All of the results presented are from at least three independent experiments and all of the data points shown indicate the mean ± standard deviation (s.d.) from n > 3 Organ Chips unless otherwise mentioned. Tests for statistically significant differences between groups were performed using one-way ANOVA followed by Tukey multiple comparison, statistical analyses were performed using GraphPad Prism 9.0.2.

## RESULTS

### Human Vagina Chip

We engineered a human Vagina Chip by co-culturing primary human vaginal epithelium on the top surface of an extracellular matrix-coated porous membrane within the top channel of a two-channel microfluidic chip with primary human uterine fibroblasts on the lower surface of the same membrane in the bottom parallel channel to recreate the vaginal epithelial-stromal interface *in vitro* (**Fig. 1A**), which has been shown to be important for development of the vaginal epithelium[35, 36]. The top and bottom channels of the Vagina Chip were respectively perfused with epithelial and stromal growth medium for 5 days to expand cell populations before replacing the epithelium medium with HBSS (LB/+G) [pH ~ 4.7] and the stromal medium with a differentiation medium that supports optimal viability and epithelial stratification (see **Methods**). The medium was continuously perfused through the lower channel and intermittently through the upper epithelial channel to mimic episodic flow of mucus through the vagina. These culture conditions resulted in spontaneous differentiation of a multilayered, stratified, squamous vaginal epithelium with a thickness of ~75-90 μm when cryosectioned along the vertical axis and stained with eosin and DAPI (**Fig. 1A**). Immunofluorescence microscopic imaging of vaginal epithelium for various tissue-specific basal, suprabasal, and superficial markers, including cytokeratin 5 (CK5), CK14, CK13, CK15, and involucrin, confirmed the presence of a well differentiated vaginal epithelium on-chip (**Fig. 1B**). The engineered vaginal epithelium also expressed cell-cell adhesion molecules that contribute to epithelial junctional complex formation, including E-cadherin, zonula occludens-1 (ZO-1), desmoglein-1 and −3 (DSG1 and DSG3) (**Fig. 1B**). The presence and absence of these proteins in different layers of vaginal epithelium on-chip recapitulated their locations observed human vagina *in vivo* (**Table 1**).

**Fig. 1.**
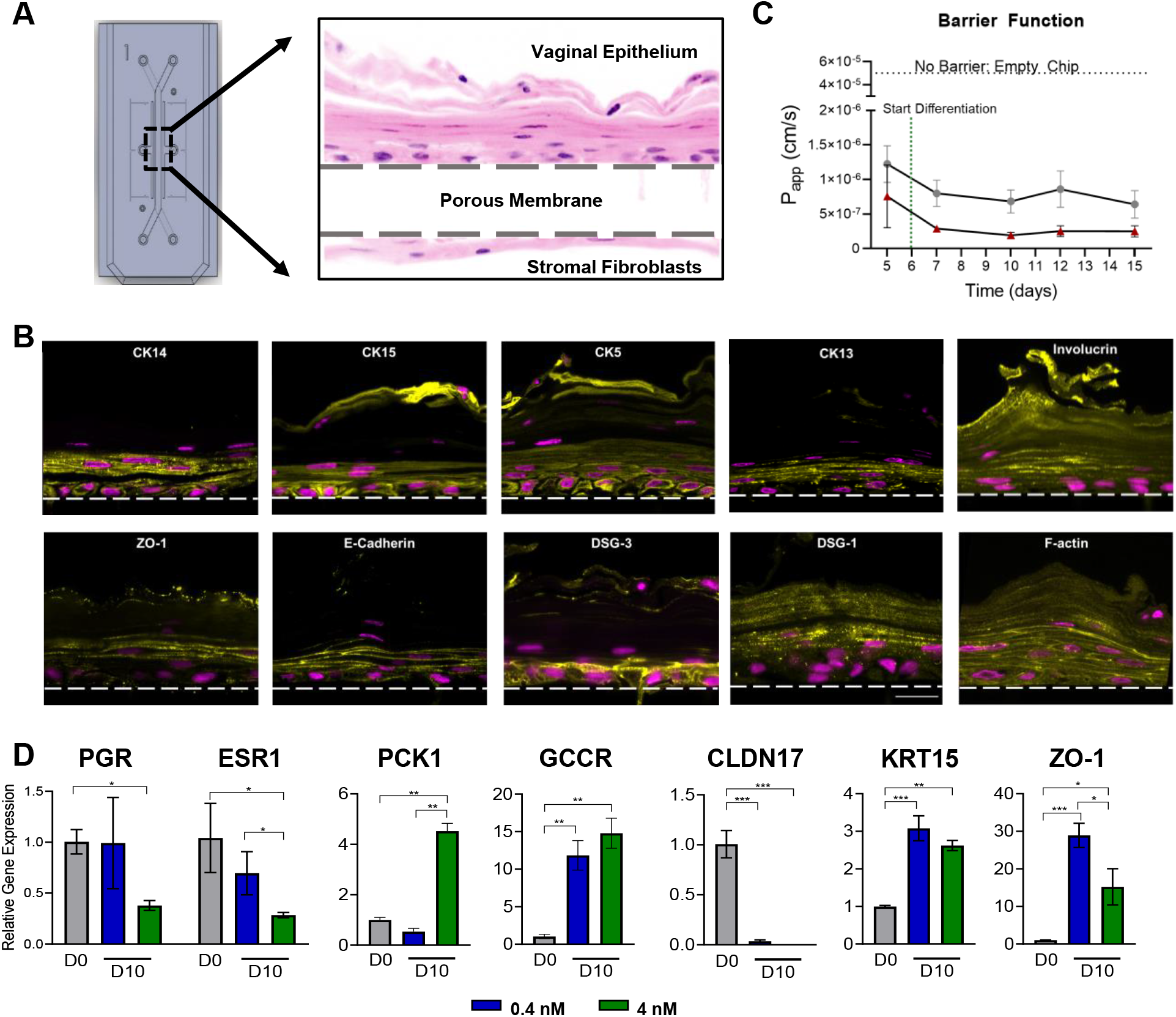
Characterization of a Microfluidic Human Vagina-on-a-Chip Model. **A**) A diagram of a two-channel Organ Chip viewed from above (left) and a higher magnification micrograph through a cross section of a Vagina Chip showing a stratified human vaginal epithelial cells cultured in the top channel atop a 50 μm thick porous membrane (dashed lines indicate top and bottom surface) with human uterine fibroblasts cultured on the opposed side of the membrane in the lower channel (right). **B)** Representative cross-sectional immunofluorescence micrographs of human Vagina Chips showing squamous stratified vaginal epithelium immunostained for CK14, CK15, CK5, CK13, Involucrin, ZO-1, E-cadherin, DSG-3, DSG-1, and F-actin (to show all cells). Dashed line, upper boundary of the porous membrane; yellow, different markers; magenta, DAPI-stained nuclei. **C)** A graph showing the changes in apparent permeability (P_app_) of the vaginal tissue barrier measured on-chip for two human donors 05328 (Hispanic) and 04033 (Caucasian) measured by quantifying Cascade Blue transport. Data are presented as mean ± s.d.; n=4. **D)** RT-qPCR results showing relative mRNA expression of ESR1, PGR, PCK1, GCGR, KRT15, CLDN17, and ZO-1 in the Vagina Chip before (D0) and after differentiation on day 10 in the presence or absence of 0.4 nM (blue) or 4 nM (green) β-estradiol (D10).

**Table 1.** Expression of various differentiation markers in different layers of vaginal epithelium on-chip

| Marker Type | Marker | Expression On-Chip | Expression in vivo | Reference |
| --- | --- | --- | --- | --- |
| Stratified Differentiation | CK13 | Suprabasal | Suprabasal | 46, 51 |
|  | CK14 | Basal | Basal | 46,47, 52 |
|  | CK15 | Basal | Basal | 48, 50 |
|  | Involucrin | Multiple cell layers | Multiple cell layers | 49,51 |
| Cell-Cell Adhesion | E-Cadherin | Multiple cell layers | Multiple cell layers | 47,49 |
|  | ZO-1 | Superficial | Superficial | 50,54,55 |
|  | Desmoglein 3 (DSG3) | Basal | Basal | 53 |
|  | Desmoglein 1 (DSG1) | Multiple cell layers<br>Greater- Superficial | Multiple cell layers<br>Greater-Superficial | 49 |

Co-culture of the vaginal epithelial cells and fibroblasts on-chip also resulted in establishment of a strong and stable epithelial permeability barrier, as measured by quantifying the apparent permeability (P_app_) using the small fluorescent biomarker, Cascade Blue (550 Da), which was sustained at 10^-6^ to 10^-7^ cm/s for up to 15 days of culture (**Fig. 1C**). Moreover, similar differentiation-induced decreases in permeability and maintenance of this high permeability barrier for at least 2 weeks of culture was observed using Vagina Chips lined with vaginal epithelial cells obtained from two donors with different ethnicity (Caucasian and Hispanic) (**Fig. 1C**). Although these are only the results of one donor each and we do not know their health status, it is interesting that chips lined with vaginal epithelial cells from the Hispanic donor appeared to form a slightly stronger barrier compared to those created with Caucasian donor cells.

This differentiation protocol was carried out in a basal growth medium containing the female sex hormone β-estradiol at a high concentration (4 nM) that mimics its peak level in blood during the human menstrual cycle *in vivo*[37]. Under these conditions, we observed down-regulation of expression of genes encoding estrogen receptor 1 (ESR1), progesterone receptor (PGR), and claudin 17 (CLDN17), while Phosphoenolpyruvate Carboxykinase 1 (PCK1), glucagon receptor (GCGR), keratin 15 (KRT15), and ZO-1 were significantly upregulated when measured using RT-qPCR on day 10 of culture compared to the pre-differentiation state (day 0) (**Fig. 1D**). Importantly, when we perfused chips with medium containing a low level of β-estradiol (0.4 nM) that mimics its nadir levels in the blood during the menstrual cycle[37], we found that the vaginal epithelium in these chips failed to significantly down regulate the PGR and ESR1 genes, as observed with the higher peak level (**Fig. 1D**). Exposure to the lower β-estradiol level was equally effective at suppressing expression of CLDN17 and inducing expression of GCGR and KRT15, however, ZO-1 expression levels appeared to be even more highly sensitive to the lower dose of estradiol (**Fig. 1D**). Thus, the Vagina Chip is able to recapitulate human vaginal epithelium responsiveness to variations in sex hormone levels *in vitro.*

### Co-culture of the Vagina Chip with optimal L. crispatus-containing microbiome

Because oxygen concentrations in the human vagina are low[38, 39] and most of the bacteria comprising the vaginal microbiota are strict or facultative anaerobes, we first simulated the O_2_ gradient generated within the Vagina Chip under the aerobic culture conditions we utilized to ensure that the environment is appropriate for microbial co-culture using a COMSOL-based two-dimensional model (**Supplementary Fig. S1A)**. The oxygen-saturated medium and diffusion of oxygen into the chip from the incubator were modeled as the main sources of O_2_ influx whereas cellular oxygen consumption was the sole source of loss. This analysis revealed that the oxygen concentration on top of the epithelial layer in the upper channel is ~0.11 mol/m^3^ (approximately 10% O_2_) (**Supplementary Fig. S1B,C)**, which is sufficiently low to support the growth of vaginal microbiota.

We tested the ability of each of the three *L. crispatus* multi-strain (OC1, OC2, and OC3) consortia to grow in the Vagina Chip and assessed their effects on host responses. These studies revealed that total live culturable bacteria could be isolated from the chip effluents daily throughout a 72 hour experiment and digested vaginal epithelial tissue after 72 hour to determine whether the Vagina Chips were co-cultured with a single *L. crispatus* strain (C0006A1) or any of the OC1, OC2 and OC3 *L. crispatus* multi-strain consortia (**Fig. 2A**). Some of the bacteria within these optimal consortia also remained adherent to the epithelium in the chip as demonstrated by quantifying the percent of inoculated bacteria that were culturable in the digested vaginal epithelial tissue on day 3 (**Fig. 2B)**.

**Fig. 2.**
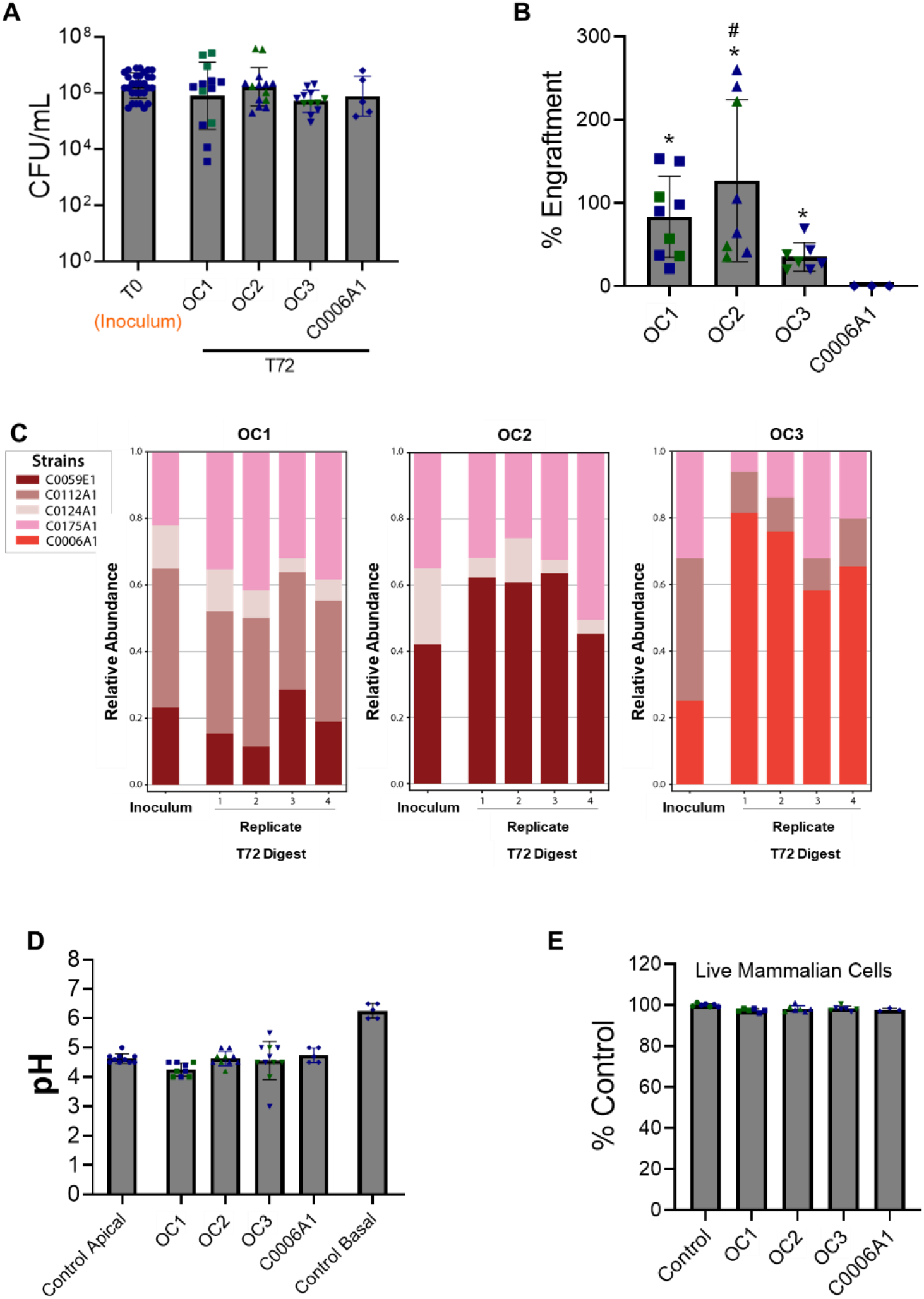
Culture of *L. crispatus* Consortia in the Vagina Chip. **A)** Total CFU/ml determined by quantifying non-adherent bacteria in effluents from the apical epithelial channel combined with counts of adherent bacteria measured within epithelial digets at 72 hours compared to the original inoculum (T0). **B)** Percent engraftment of OC1, OC2, OC3 and C0006A1 consortia bacteria in Vagina Chips calculated by quantifying the bacteria recovered from the chip (digest) 72 hours post inoculation compared to the T0 inoculum. Significance was calculated by one-way ANOVA; *, P < 0.05 vs. C0006A1, #, P < 0.05 vs. OC1. **C)** The pH values measured in the medium within the apical channel of the Vagina Chip cultured in the absence or presence of the OC1, OC2, OC3 and C0006A1 consortia at 72 hours post inoculation compared to pH measured in the basal channel. **D)** Percent viability of vaginal epithelial cells assessed by calculating the number of live cells relative to control using trypan blue exclusion assay. In **A-D,** each data point indicates one chip; different colored points indicate chips from different donors; data are presented as mean±s.d. In the lower graphs, results of metagenomics-based strain ratio analyses of the engrafted *L. crispatus* OC1 (**E**), OC2 (**F**), and OC3 (**G**) consortia present within epithelial digests relative to the original inoculum are shown after 72 hours of direct culture with the vaginal epithelium on-chip. 1-4 indicate results from 4 different chips.

We then carried out metagenomics-based strain ratio analyses of the engrafted *L. crispatus* consortia to assess the degree of cooperativity among the component bacterial strains. For all 3 multi-strain consortia, we detected the presence of all strains on chips after 72 hours of direct contact with the vaginal epithelium (**Fig. 2C**). A similar ratio of different strains of engrafted OC1 and OC2 consortia was observed when compared with their respective inoculums, but the C0006A1 strain became more predominant in the OC3 consortia (~25% to ~70%). This is interesting given that the C0006A1 did not engraft in the chip when cultured alone (i.e., not as part of a multi-strain consortium)(**Fig. 2B**). The strain-level stability of different *L. crispatus* consortia also appeared to remain relatively constant in the Vagina Chips when four different replicates were compared (**Fig. 2C**).

### Maintenance of physiological pH

The epithelium of the Vagina Chip was cultured in an HBSS solution (pH ~4.7) to mimic the physiological pH experienced by vaginal epithelium *in vivo*, and the chip was able to maintain this pH (**Fig. 2D**) as well as epithelial cell viability (**Fig. 2E**), when cultured in the presence and absence of *L. crispatus* bacteria, either as a single strain (C0006A1) or within the OC1, OC2, or OC3 consortia.

### Lactate production in the Vagina Chip

The D- and L-enantiomers of lactic acid both have antimicrobial effects[3, 4], however, while vaginal epithelium can only produce L-lactic acid, *L. crispatus* has the ability to produce both isomers, making D-lactic acid a biomarker for metabolically active *L. crispatus* bacteria[40]. As expected, we detected L-lactate in all Vagina Chips containing human vaginal epithelial cells, and this was the only isomer present in control chips and in those inoculated with *L. crispatus* strain C0006A1 which failed to engraft on the vaginal epithelium in our experiments (**Fig. 3A**). In contrast, both L- and D-lactate were detected in Vagina Chips containing OC1, OC2 and OC3 microbial consortia after 72 hours, although only the chips containing the OC2 and OC3 consortia exhibited levels (0.33 mM and 0.29 mM, respectively) similar to those observed *in vivo* (0.32 mM)[41] (**Fig. 3B**).

**Fig. 3.**
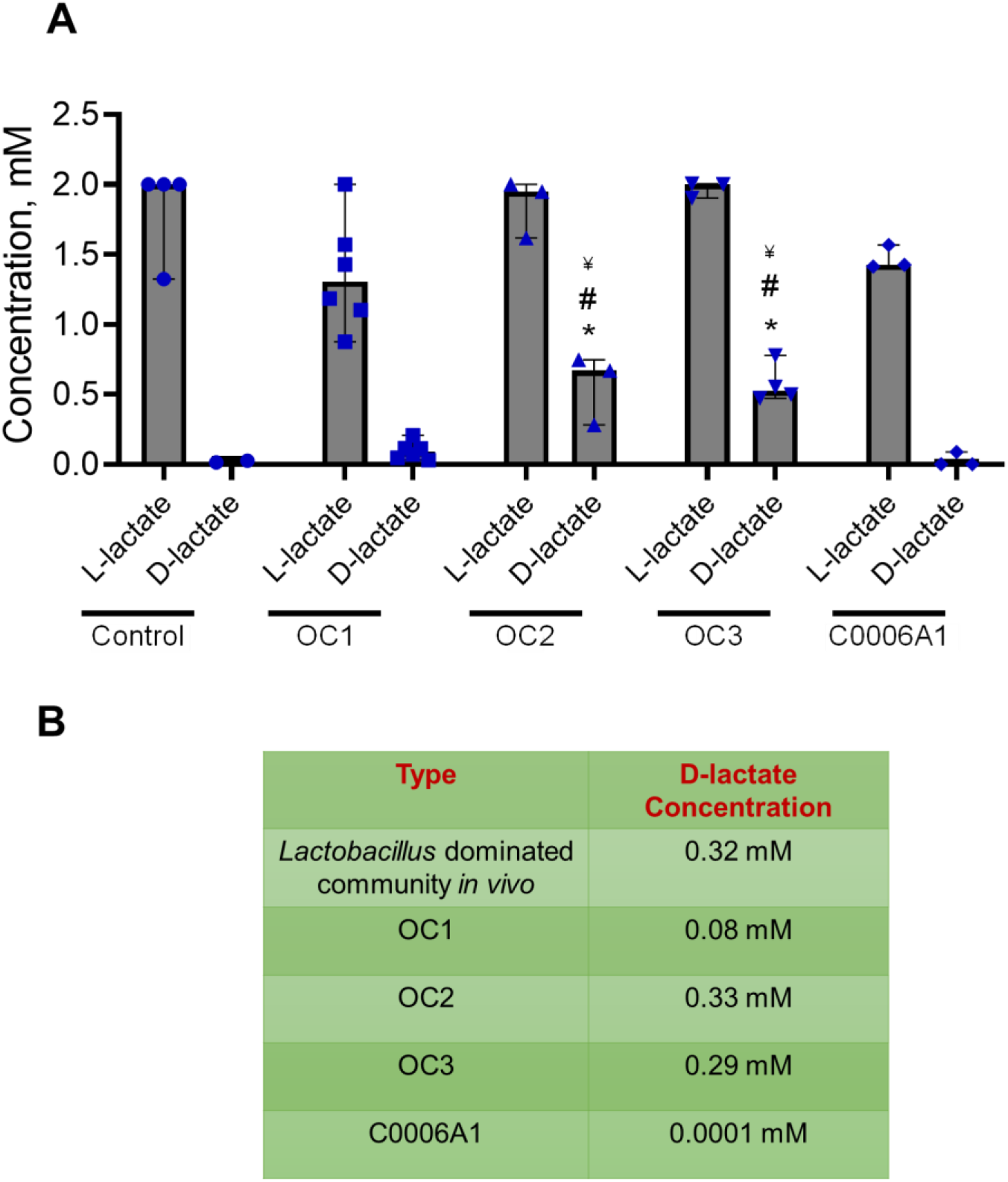
D-lactate Production in Vagina Chip. **A)** D-lactate and L-lactate levels measured in the effluent of the epithelial channel of Vagina Chip cultured in the absence (control) or presence single strain (C0006A1) or multi-strain (OC1, OC2, OC3) *L. crispatus* consortia at 72 hours post inoculation. Each data point indicates one chip; data are presented as median with 95% CI. Significance was calculated by one-way ANOVA; *, P < 0.05 vs. control, #, P < 0.05 vs. OC1, ^¥^, P < 0.05 vs. C0006A1. **B)** Table showing median D-lactate concentrations (mM) measured in the vagina of women with *Lactobacillus*-dominated communities compared to concentrations measured in Vagina Chips cultured with different *L. crispatus* consortia (OC1, OC2, OC3) and the C0006A1 strain.

### Modulation of innate immune responses by optimal vaginal microbiota

In the vagina, *Lactobacillus* species are believed to provide benefit by suppressing inflammation[3, 42, 43]. Consistent with this observation, we found that when the vaginal epithelial cells were grown on-chip with or without the OC1, OC2, or OC3 consortia, or *L. crispatus* strain C0006A1, we observed a statistically significant downregulation of multiple proinflammatory cytokines, including interleukin-6 (IL-6), IL-8, IL-1α, IL-1β, and interferon-γ inducible protein-10 (IP-10) after 72 hours of co-culture compared to controls Vagina Chips **(Fig. 4)**. These results with the Vagina Chip demonstrate that *L-crispatus* containing consortia can directly influence the epithelium to dampen production of inflammatory cytokines, even in the absence of immune cells.

**Fig 4.**
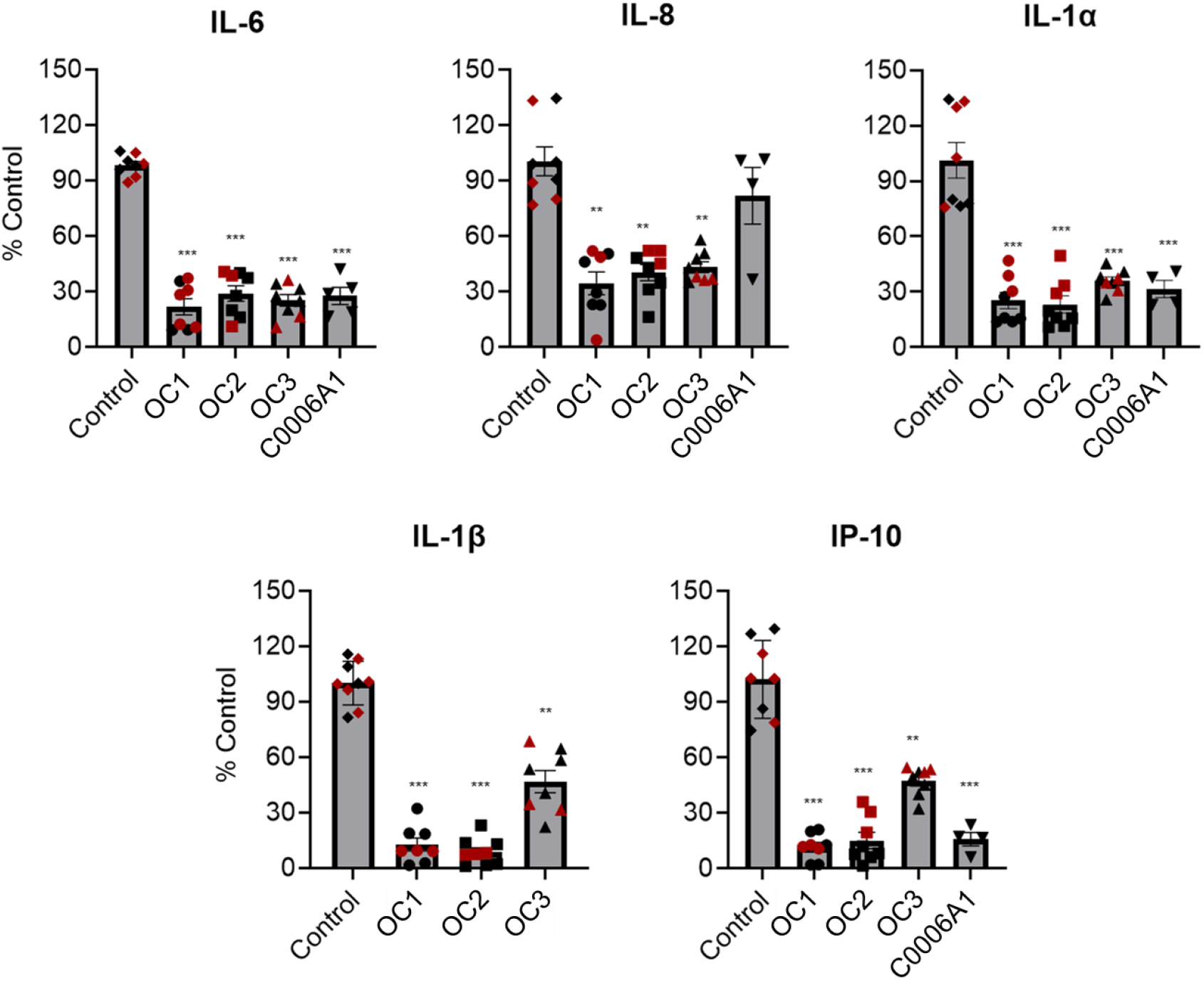
Suppression of the innate immune response by *L. crispatus* containing consortia in the Vagina Chip. The levels of cytokines (IL-6, IL-8, IL-1α, IL-1β, and IP-10) measured in effluents of Vagina Chips cultured with OC1, OC2, OC3 and C0006A1 consortia are show relative to control chips without bacteria. Each data point indicates one chip; different colored points indicate chips from different donors. Data are presented as mean ± sem; significance was calculated by one-way ANOVA; ***, P< 0.0001; **, P < 0.001.

### Culture of non-optimal G. vaginalis containing vaginal microbiome in the Vagina Chip

We also studied the effects of co-culturing non-optimal vaginal microbiome in the Vagina Chip by inoculating the chips (~10^6^ CFU/chip) with consortia containing either G. *vaginalis E2* and *E4* combined with *P. bivia BHK8* and *A. vaginae* (BVC1) or only the two G. *vaginalis strains*(BVC2) on day 11 of differentiation. Quantification of total bacterial count by cultivation indicated that members of both BVC1 and BVC2 consortia remained present and viable on-chip throughout this 3 day study, although the strain ratios were not determined (**Fig. 5A)**. Based on the CFU/mL of digested epithelium, we observed that both consortia were able to colonize the vaginal tissue and thrive on the Vagina Chip, as the total CFU/ml measured in the epithelial digests plus the effluents increased over the 3 day culture from 10^6^ to ~10^8^ CFU/ml (**Fig. 5A**). Co-culture of the BVC1 consortium on-chip resulted in a physiologically relevant and statistically significant increase in pH to ~5.1, while no pH change (~4.7) was observed in presence of BVC2 (**Fig. 5B**). We also observed a reduction in vaginal epithelial cell viability when cultured in the presence of either the BVC1 or BVC2 consortium (**Fig. 5C**). As expected, no D-lactate was detected in Vagina Chips containing BVC1 or BVC2 consortia (not shown). Importantly, in contrast to the *L. crispatus* containing consortium both *G. vaginalis* containing consortia induced statistically significant increases in the production of multiple proinflammatory cytokines (IL-6, IL-8, IL-1β and IP-10) after 72 hours of co-culture (**Fig. 5D versus Fig. 4**), similar to *in vivo* observations[44, 45].

**Fig. 5.**
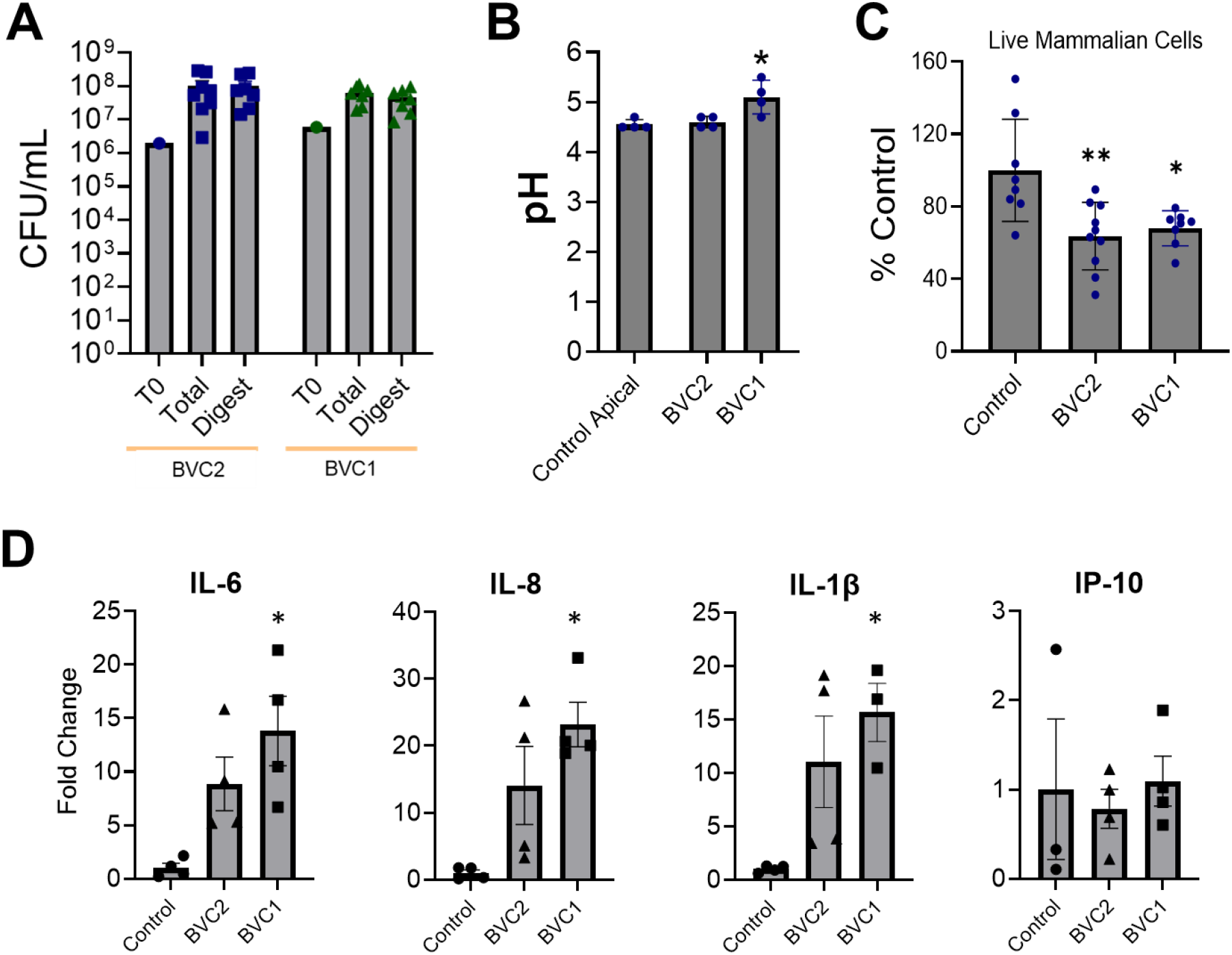
Culture of non-optimal *G. vaginalis* containing Consortia in the Vagina Chip. **A**) Total CFU/ml of BVC1 and BVC2 consortia bacteria measured in effluents from the vaginal epithelium-lined channel at 24, 48, and 72 hours and epithelial tissue digest at 72 hours relative to the starting inoculum (T0). Total, total CFUs measured in effluents + digest; Digest, CFU in the tissue digest. **B)** pH measured in medium in the epithelial channel of Vagina Chips cultured in the presence or absence of *G. vaginalis* containing and BVC1 and BVC2 consortia for 72 hours. **C)** Percent viability of vaginal epithelial cells cultured on-chip in the presence of BVC1 or BVC2 consortia assessed by calculating the number of live cells relative to control using a trypan blue exclusion assay. **D)** The levels of cytokines (IL-6, IL-8, IL-1α, IL-1β, and IP-10) measured in effluents of Vagina Chips cultured with BVC1 and BVC2 consortia are shown relative to control chips. Each data point indicates one chip; data are presented as mean ±sem; significance was calculated by one-way ANOVA; *, P < 0.01,**, P < 0.001.

## DISCUSSION

In this study, we set out to explore whether Organ Chip technology can be used to develop a preclinical model of human vagina-microbiome interactions, which could potentially be used for discovery and assessment of potential microbiome-based therapeutics. The microfluidic Vagina Chip lined by primary human vaginal epithelial cells interfaced with uterine fibroblasts that we engineered forms a squamous stratified vaginal epithelium expressing various differentiation markers in correct locations that closely mimic those observed of human vaginal epithelium *in vivo* [46–55]. The Vagina Chip also exhibits a tight tissue permeability barrier, maintains a physiologically relevant low pH, responds to estrogen hormone, and creates an oxygen gradient that enables stable co-culture with microbial communities including both optimal *L. crispatus* strain containing consortia and non-optimal *G. vaginalis* strain containing consortia. The Vagina Chip was used to study host-microbiome interactions using a single strain of *L. crispatus* as well as three different multi-strain *L. crispatus* consortia, which resulted in D-lactate accumulation and suppression of inflammatory cytokine production on-chip, thus mimicking their beneficial effects on vaginal health observed *in vivo.* In contrast, when *G. vaginalis* or mixed consortia containing pathogen G. vaginalis and other anaerobes commonly found in non-optimal vaginal microbiota were cultured on-chip, secretion of inflammatory cytokines and chemokines increased and this was accompanied by vaginal cell injury. Interestingly, we also observed a single *L. crispatus* strain (C0006A1) failed to engraft in the Vagina Chip even though the same strain thrived on-chip when it was co-cultured as part of the multi-strain OC3 consortium. These findings are consistent with past work which suggests that colonization or engraftment of *L. crispatus* consortia in the vagina may result in enduring changes to the total microbiome composition and consequently helping to prevent recurrent vaginal dysbiosis[56].

Past studies have shown short-term adhesion of lactobacilli (especially *L. crispatus*) to cell monolayer cultures and engraftment in animal models, although live bacterial cell numbers (CFU) were not quantified[57, 58]. Monolayer cultures lack key structural and functional features of living three-dimensional tissues that are important to mimic host-microbiome interactions, such as the presence of a multi-layered. The immunomodulatory properties of *L. crispatus* bacteria have been demonstrated previously using cervicovaginal monolayer cultures or transwell models[42, 58] or 3D aggregrates[59] however, some of these studies used immortalized cells[42, 58] or human cells derived from ectocervical tissue or vulval epidermoid carcinoma[46, 60] rather than healthy vaginal epithelium as we did in the present study. In addition, typically these static culture systems could only be used for short-term (<24 hr) studies due to bacterial overgrowth. In contrast, our ability to reconstitute the vaginal epithelial-stromal interface and expose both compartments to dynamic fluid flow independently enabled longer term (> 3 day) co-culture of microbiome in direct contact with living human vaginal epithelium.

The Vagina Chip also expresses multiple structural and functional markers that mimic those observed *in vivo*, which are critical for support of a living microbiome and maintenance of vaginal health[46–55]. For example, PCK1 is a component of the molecular machinery involved in production of glycogen, which is thought to be the key nutrient for vaginal *lactobacilli* [61, 62]. Accumulation of glycogen and thickening of the vaginal epithelium are also induced by increased estrogen levels in humans[63] as well as animal studies,[64, 65] and this is consistent with our observation that high levels of estrogen upregulated genes involved in gluconeogenesis and glycogen synthesis and downregulate estrogen receptor genes in the Vagina Chip. Moreover, the O_2_ partial pressure in the human vagina is within the hypoxic range which supports vaginal lactobacilli production of D-lactate[39, 66] and we experimentally confirmed that this occurs on-chip as well. Thus, the human Vagina Chip offers a more physiologically relevant and versatile experimental system for *in vitro* studies on host-microbiome interactions than existing *in vitro* models.

*L. crispatus* produce D- and L-lactate, which have antimicrobial and immunomodulatory properties that help to maintain an optimal *Lactobacillus*-dominant community[4, 63]. For example, D-lactate blocks chlamydia infection *in vitro*[67, 68], and inhibits Toll-like receptor (TLR) agonist-elicited production of inflammatory mediators in study using an epithelial cell line[69]. Lactic acid decreases production of pro-inflammatory mediators (IL-6 and IL-8) in cultured cervical epithelium[69] and lactate production by *L. crispatus* and *L. gasseri* has been shown to prevent infection by *Chlamydia trachomatis,*[68] suppress growth of *Escherichia coli*[70], and *Neisseria gonorrhoeae* bacteria[71]. Importantly, we observed physiological levels of D-lactate on-chip [41] when the vaginal epithelium was co-cultured with multi-strain OC2 and OC3 consortia, whereas levels were much lower with the OC1 consortium. Thus, the human Vagina Chip may be useful for assessing relative efficacy of different live biotherapeutic product formulations in terms of their ability to produce D-lactate at the surface of the vaginal epithelium, and hence, suppress pathogen infection and inflammation.

In addition to acting as a physical barrier to infections, vaginal epithelial cells generate a powerful innate immune response to non-optimal bacteria associated with BV by producing inflammatory cytokines and anti-microbial products, such as defensins and lysozymes [43, 72], while a microbiome dominated by *L. crispatus* that is considered optimal reduces the pro-inflammatory response as shown, for example, in cervicovaginal cell cultures stimulated with various TLR agonists[3, 42]. Vaginal epithelial cells co-cultured with *Lactobacillus* isolates from women with optimal microbiome communities also produce lower levels of pro-inflammatory cytokines than isolates from non-optimal microbial communities[58]. Downregulation of multiple pro-inflammatory cytokines and chemokines was observed in the present study when vaginal epithelial cells were co-cultured with multi-strain *L. crispatus* consortia in the Vagina Chip. Conversely, several of these cytokines are upregulated in clinical samples with non-optimal microbiota associated with preterm birth[73]. In contrast, when we cultured Vagina Chips with either *G. vaginalis* strains alone or as part of consortia containing these potentially pathogenic strains, we observed epithelial cell injury and significant upregulation of these same proinflammatory molecules. These findings highlight the ability of the Vagina Chip to recapitulate host-vaginal microbiome interactions that are observed *in vivo* which play a central role in vaginal health and to discriminate between probiotic and dysbiotic bacterial consortia *in vitro*.

BV, which is the most common vaginal condition in reproductive-aged women, is characterized by increased vaginal discharge and changes in the vaginal microbiota. During BV, beneficial *Lactobacilli* are displaced by an array of strict and facultative anaerobes, including *Gardnerella, Prevotella, Mobiluncus,* and *Atopobium* species[74]. Given the unsatisfactory efficacy of the current treatment regime to prevent recurrent dysbiosis[56], the use of *L. crispatus-based* therapeutic strategies is gaining interest. For example, the first living *L. crispatus* probiotic therapeutic product that was derived from a human vaginal microbiome sample (LACTIN-V) showed promising engraftment in the clinical trial with 79% of participants showing qPCR detection of LACTIN-V bacteria after 12 weeks[56]. However, the development of new and even more effective BV therapeutics, including live probiotic therapies, would benefit from the availability of human relevant preclinical models that also enable assessment of the effects of dynamic host-microbiome interactions. Current approaches that are used to study of interactions between human vagina and healthy or dysbiotic microbiome, as well as to develop live biotherapeutics, utilize animal or *in vitro* models for preclinical analysis. But different species have distinct microbiomes and some of the existing *in vitro* human cell culture models are also limited in terms of their physiological mimicry and their ability to support extended co-culture studies with living bacteria.

Most importantly, our demonstration that the human Vagina Chip can be used to investigate human vagina-microbiome interactions using single- and multi-strain consortia containing *L. crispatus* strains as well as dysbiotic *Gardnerella-containing* bacterial strains, suggests that it could be used as a new preclinical model to advance therapeutic development in the future as there is no other way to assess these activities *in vitro*. Various species and strains of *Lactobacillus* (e.g., *L. crispatus, L. gasseri, L. acidophilus, L. fermentum, L. rhamnosus)* have been assessed as potential probiotics for the treatment of vaginal dysbiosis and specifically BV[75, 76]. We chose to explore the effects of *L. crispatus* strains on vaginal tissue in our Organ Chip model because they are highly associated with positive gynecological outcomes and it is the dominant *Lactobacillus* species in healthy vaginal microbiomes[75]. Consistent with these observations, our Vagina Chip results clearly show that vaginal epithelial cells remain healthy and viable when in direct contact with either a single *L. crispatus* strain or a multi-strain consortium, although the single strain failed to successfully engraft on-chip. In contrast, culture with a potential vaginal pathogen, *G. vaginalis* (either alone or as part of a more complex microbial consortium) resulted in epithelial injury and enhanced inflammation.

In summary, the human Vagina Chip supports spontaneous differentiation of squamous stratified vaginal epithelium, forms a strong barrier, responds to hormones, and generates a microbiome supporting oxygen-gradient. In addition, we demonstrated that multi-strain *L. crispatus* consortia outperform single-strain *L. crispatus* in terms of engraftment, D-lactate production and suppression of an innate immune response. In contrast, when dysbiosis associated *G. vaginalis* containing microbiota were cultured in the Vagina Chip, epithelial injury and enhanced inflammation resulted. Taken together, these data indicate that the human Vagina Chip offers a new model to study host-vaginal microbiome interactions in both optimal and non-optimal states, as well as providing a human relevant preclinical model for development and testing of reproductive therapeutics, including live bio-therapeutics products for BV.

## AUTHORS CONTRIBUTIONS

G.M. designed *in vitro* experiments with the help of D.E.I., R.P.B, E.D., T.T., A.S., J.R., G.G., I. H-P., and S.R-N.. G.M, E.D and T.T performed *in vitro* experiments and analyzed the data, working with D.E.I, who also supervised all the work. A.S, R.P, S.S, J.G., J.R. and V.H. assisted in performing experiments and analyzing the data. G.M, E.D, T.T., J.R. and D.E.I. wrote the manuscript.

## ACKNOWLEDGMENTS

This research was sponsored by the funding from the Bill and Melinda Gates Foundation (OPP1173198 & INV-035977 to D.E.I., OPP1189217 to J.R. and and INV-031642 to S.R-N.) and the Wyss Institute for Biologically Inspired Engineering (D.E.I.). The authors would like to thank Vadika Mishra and Dr. Sasan Jalili-Firoozinezhad for their assistance during the early exploratory phase of this project, and Dr. Amir Bein for helpful discussions.

## REFERENCES

1. Proctor LM, Creasy HH, Fettweis JM, Lloyd-Price J, Mahurkar A, Zhou W, et al. The Integrative Human Microbiome Project. Nature. 2019;569:641–8.

2. Ravel J, Gajer P, Abdo Z, Schneider GM, Koenig SSK, McCulle SL, et al. Vaginal microbiome of reproductive-age women. Proceedings of the National Academy of Sciences. 2011;108 Supplement 1:4680 LP – 4687.

3. Delgado-Diaz DJ, Tyssen D, Hayward JA, Gugasyan R, Hearps AC, Tachedjian G. Distinct Immune Responses Elicited From Cervicovaginal Epithelial Cells by Lactic Acid and Short Chain Fatty Acids Associated With Optimal and Non-optimal Vaginal Microbiota. Frontiers in cellular and infection microbiology. 2020;9:446.

4. Tachedjian G, Aldunate M, Bradshaw CS, Cone RA. The role of lactic acid production by probiotic Lactobacillus species in vaginal health. Research in Microbiology. 2017;168:782–92.

5. Stoyancheva G, Marzotto M, Dellaglio F, Torriani S. Bacteriocin production and gene sequencing analysis from vaginal Lactobacillus strains. Archives of Microbiology. 2014;196:645–53.

6. Zheng J, Gänzle MG, Lin XB, Ruan L, Sun M. Diversity and dynamics of bacteriocins from human microbiome. Environmental Microbiology. 2015;17:2133–43.

7. van de Wijgert JHHM, Jespers V. The global health impact of vaginal dysbiosis. Research in Microbiology. 2017;168:859–64.

8. Juliana NCA, Suiters MJM, Al-Nasiry S, Morré SA, Peters RPH, Ambrosino E. The Association Between Vaginal Microbiota Dysbiosis, Bacterial Vaginosis, and Aerobic Vaginitis, and Adverse Pregnancy Outcomes of Women Living in Sub-Saharan Africa: A Systematic Review. Frontiers in Public Health. 2020;8:850.

9. Janulaitiene M, Paliulyte V, Grinceviciene S, Zakareviciene J, Vladisauskiene A, Marcinkute A, et al. Prevalence and distribution of Gardnerella vaginalis subgroups in women with and without bacterial vaginosis. BMC Infectious Diseases. 2017;17:394.

10. Eastment MC, McClelland RS. Vaginal microbiota and susceptibility to HIV. AIDS (London, England). 2018;32:687–98.

11. Brunham RC, Gottlieb SL, Paavonen J. Pelvic Inflammatory Disease. New England Journal of Medicine. 2015;372:2039–48.

12. van de Wijgert JHHM, Verwijs MC, Gill AC, Borgdorff H, van der Veer C, Mayaud P. Pathobionts in the Vaginal Microbiota: Individual Participant Data Meta-Analysis of Three Sequencing Studies. Frontiers in Cellular and Infection Microbiology. 2020;10:129.

13. Brown RG, Marchesi JR, Lee YS, Smith A, Lehne B, Kindinger LM, et al. Vaginal dysbiosis increases risk of preterm fetal membrane rupture, neonatal sepsis and is exacerbated by erythromycin. BMC medicine. 2018;16:9.

14. Fettweis JM, Serrano MG, Brooks JP, Edwards DJ, Girerd PH, Parikh HI, et al. The vaginal microbiome and preterm birth. Nature Medicine. 2019;25:1012–21.

15. Lagenaur LA, Hemmerling A, Chiu C, Miller S, Lee PP, Cohen CR, et al. Connecting the Dots: Translating the Vaginal Microbiome Into a Drug. The Journal of Infectious Diseases. 2021;223 Supplement_3:S296–306.

16. Schenk M, Grumet L, Sternat J, Reinschissler N, Weiss G. Effect of probiotics on vaginal Ureaplasma parvum in women suffering from unexplained infertility. Reproductive BioMedicine Online. 2021. https://doi.org/doi.org/10.1016/j.rbmo.2021.06.004.

17. Miller E, Beasley D, Dunn R, Archie E. Lactobacilli Dominance and Vaginal pH: Why is the Human Vaginal Microbiome Unique? Frontiers in Microbiology. 2016;7.

18. Kim HJ, Huh D, Hamilton G, Ingber DE. Human gut-on-a-chip inhabited by microbial flora that experiences intestinal peristalsis-like motions and flow. Lab on a chip. 2012;12:2165–74.

19. Jalili-Firoozinezhad S, Gazzaniga FS, Calamari EL, Camacho DM, Fadel CW, Bein A, et al. A complex human gut microbiome cultured in an anaerobic intestine-on-a-chip. Nature Biomedical Engineering. 2019;3:520–31.

20. Kim HJ, Li H, Collins JJ, Ingber DE. Contributions of microbiome and mechanical deformation to intestinal bacterial overgrowth and inflammation in a human gut-on-a-chip. Proceedings of the National Academy of Sciences. 2016;113:E7 LP–E15.

21. Ingber DE. Is it Time for Reviewer 3 to Request Human Organ Chip Experiments Instead of Animal Validation Studies? Advanced Science. 2020;7:2002030.

22. Huh D, Matthews BD, Mammoto A, Montoya-Zavala M, Yuan Hsin H, Ingber DE. Reconstituting organ-level lung functions on a chip. Science. 2010;328:1662–8.

23. Bhatia SN, Ingber DE. Microfluidic organs-on-chips. Nature Biotechnology. 2014;32:760–72.

24. Elfer KN, Sholl AB, Wang M, Tulman DB, Mandava SH, Lee BR, et al. DRAQ5 and Eosin (‘D&E’) as an Analog to Hematoxylin and Eosin for Rapid Fluorescence Histology of Fresh Tissues. PLOS ONE. 2016;11:e0165530.

25. Gazzaniga FS, Camacho DM, Wu M, Silva Palazzo MF, Dinis ALM, Grafton FN, et al. Harnessing Colon Chip Technology to Identify Commensal Bacteria That Promote Host Tolerance to Infection. Frontiers in Cellular and Infection Microbiology. 2021;11:105.

26. Ma B, France MT, Crabtree J, Holm JB, Humphrys MS, Brotman RM, et al. A comprehensive non-redundant gene catalog reveals extensive within-community intraspecies diversity in the human vagina. Nature Communications. 2020;11:940.

27. Ravel J, Brotman RM, Gajer P, Ma B, Nandy M, Fadrosh DW, et al. Daily temporal dynamics of vaginal microbiota before, during and after episodes of bacterial vaginosis. Microbiome. 2013;1:29.

28. Bloom SM, Mafunda NA, Woolston BM, Hayward MR, Frempong JF, Abai AB, et al. Cysteine dependence of Lactobacillus iners is a potential therapeutic target for vaginal microbiota modulation. Nature Microbiology. 2022;7:434–50.

29. Rotmistrovsky K & ARBmt. Best Match Tagger for Removing Human Reads from Metagenomics Datasets (NCBI/NLM, National Institutes of Health, 2011).

30. Kopylova E, Noé L, Touzet H. SortMeRNA: fast and accurate filtering of ribosomal RNAs in metatranscriptomic data. Bioinformatics. 2012;28:3211–7.

31. Chen S, Zhou Y, Chen Y, Gu J. fastp: an ultra-fast all-in-one FASTQ preprocessor. Bioinformatics. 2018;34:i884–90.

32. Li L, Stoeckert Jr CJ, Roos DS. OrthoMCL: identification of ortholog groups for eukaryotic genomes. Genome research. 2003;13:2178–89.

33. Langmead B, Salzberg SL. Fast gapped-read alignment with Bowtie 2. Nature methods. 2012;9:357–9.

34. Li H, Handsaker B, Wysoker A, Fennell T, Ruan J, Homer N, et al. The Sequence Alignment/Map format and SAMtools. Bioinformatics (Oxford, England). 2009;25:2078–9.

35. Ogawa-Tominaga M, Umezu T, Nakajima T, Tomooka Y. Stratification of mouse vaginal epithelium. 1. Development of three-dimensional models in vitro with clonal cell lines. Biology of Reproduction. 2018;99:718–26.

36. Cunha GR, Robboy SJ, Kurita T, Isaacson D, Shen J, Cao M, et al. Development of the human female reproductive tract. Differentiation. 2018;103:46–65.

37. Marsh EE, Shaw ND, Klingman KM, Tiamfook-Morgan TO, Yialamas MA, Sluss PM, et al. Estrogen levels are higher across the menstrual cycle in African-American women compared with Caucasian women. The Journal of clinical endocrinology and metabolism. 2011;96:3199–206.

38. Wagner G, Bohr L, Wagner P, Petersen LN. Tampon-induced changes in vaginal oxygen and carbon dioxide tensions. American journal of obstetrics and gynecology. 1984;148:147–50.

39. Hill DR, Brunner ME, Schmitz DC, Davis CC, Flood JA, Schlievert PM, et al. In vivo assessment of human vaginal oxygen and carbon dioxide levels during and post menses. Journal of Applied Physiology. 2005;99:1582–91.

40. Boskey ER, Cone RA, Whaley KJ, Moench TR. Origins of vaginal acidity: high D/L lactate ratio is consistent with bacteria being the primary source. 2001.

41. S. WS, Helena M-S, M. LI, Aswathi J, J. LW, J. FL, et al. Influence of Vaginal Bacteria and d- and l-Lactic Acid Isomers on Vaginal Extracellular Matrix Metalloproteinase Inducer: Implications for Protection against Upper Genital Tract Infections. mBio. 2021;4:e00460–13.

42. Rose II WA, McGowin CL, Spagnuolo RA, Eaves-Pyles TD, Popov VL, Pyles RB. Commensal Bacteria Modulate Innate Immune Responses of Vaginal Epithelial Cell Multilayer Cultures. PLOS ONE. 2012;7:e32728.

43. Pivarcsi A, Nagy I, Koreck A, Kis K, Kenderessy-Szabo A, Szell M, et al. Microbial compounds induce the expression of pro-inflammatory cytokines, chemokines and human beta-defensin-2 in vaginal epithelial cells. Microbes and infection. 2005;7:1117–27.

44. Hemalatha R, Ramalaxmi BA, KrishnaSwetha G, Kumar PU, Rao DM, Balakrishna N, et al. Cervicovaginal Inflammatory Cytokines and Sphingomyelinase in Women With and Without Bacterial Vaginosis. The American Journal of the Medical Sciences. 2012;344:35–9.

45. Morrill S, Gilbert NM, Lewis AL. Gardnerella vaginalis as a Cause of Bacterial Vaginosis: Appraisal of the Evidence From in vivo Models. Frontiers in Cellular and Infection Microbiology. 2020;10.

46. Ayehunie S, Cannon C, Lamore S, Kubilus J, Anderson DJ, Pudney J, et al. Organotypic human vaginal-ectocervical tissue model for irritation studies of spermicides, microbicides, and feminine-care products. Toxicology in Vitro. 2006;20:689–98.

47. Zhu Y, Yang Y, Guo J, Dai Y, Ye L, Qiu J, et al. Ex vivo 2D and 3D HSV-2 infection model using human normal vaginal epithelial cells. Oncotarget. 2017;8:15267–82.

48. Smedts F, Ramaekers F, Leube RE, Keijser K, Link M, Vooijs P. Expression of keratins 1, 6, 15, 16, and 20 in normal cervical epithelium, squamous metaplasia, cervical intraepithelial neoplasia, and cervical carcinoma. The American journal of pathology. 1993;142:403–12.

49. Dinh MH, Okocha EA, Koons A, Veazey RS, Hope TJ. Expression of structural proteins in human female and male genital epithelia and implications for sexually transmitted infections. Biology of Reproduction. 2012;86:1–6.

50. Yoshida S, Yasuda M, Miyashita H, Ogawa Y, Yoshida T, Matsuzaki Y, et al. Generation of stratified squamous epithelial progenitor cells from mouse induced pluripotent stem cells. PLoS ONE. 2011;6.

51. Fichorova RN, Rheinwald JG, Anderson DJ. Generation of papillomavirus-immortalized cell lines from normal human ectocervical, endocervical, and vaginal epithelium that maintain expression of tissue-specific differentiation proteins. Biology of reproduction. 1997;57:847–55.

52. Rajan N, Pruden DL, Kaznari H, Cao Q, Anderson BE, Duncan JL, et al. Characterization of an immortalized human vaginal epithelial cell line. 2000.

53. Fujii E, Funahashi S, Taniguchi K, Kawai S, Nakano K, Kato A, et al. Tissue-specific effects of an anti-desmoglein-3 ADCC antibody due to expression of the target antigen and physiological characteristics of the mouse vagina. Journal of toxicologic pathology. 2020;33:67–76.

54. Blaskewicz CD, Pudney J, Anderson DJ. Structure and function of intercellular junctions in human cervical and vaginal mucosal epithelia. Biology of Reproduction. 2011;85:97–104.

55. Oh K-J, Lee H-S, Ahn K, Park K. Estrogen Modulates Expression of Tight Junction Proteins in Rat Vagina. BioMed research international. 2016;2016:4394702.

56. Cohen CR, Wierzbicki MR, French AL, Morris S, Newmann S, Reno H, et al. Randomized Trial of Lactin-V to Prevent Recurrence of Bacterial Vaginosis. New England Journal of Medicine. 2020;382:1906–15.

57. Wang G, Zhang M, Zhao J, Xia Y, Lai PF-H, Ai L. A Surface Protein From Lactobacillus plantarum Increases the Adhesion of Lactobacillus Strains to Human Epithelial Cells. Frontiers in Microbiology. 2018;9:2858.

58. Manhanzva MT, Abrahams AG, Gamieldien H, Froissart R, Jaspan H, Jaumdally SZ, et al. Inflammatory and antimicrobial properties differ between vaginal Lactobacillus isolates from South African women with non-optimal versus optimal microbiota. Scientific Reports. 2020;10:6196.

59. Doerflinger SY, Throop AL, Herbst-Kralovetz MM. Bacteria in the Vaginal Microbiome Alter the Innate Immune Response and Barrier Properties of the Human Vaginal Epithelia in a Species-Specific Manner. The Journal of Infectious Diseases. 2014;209:1989–99.

60. Costin G-E, Raabe HA, Priston R, Evans E, Curren RD. Vaginal Irritation Models: The Current Status of Available Alternative and In Vitro Tests. Alternatives to Laboratory Animals. 2011;39:317–37.

61. Mirmonsef P, Hotton AL, Gilbert D, Burgad D, Landay A, Weber KM, et al. Free Glycogen in Vaginal Fluids Is Associated with Lactobacillus Colonization and Low Vaginal pH. PLOS ONE. 2014;9:e102467.

62. Nunn KL, Forney LJ. Unraveling the Dynamics of the Human Vaginal Microbiome. The Yale journal of biology and medicine. 2016;89:331–7.

63. Amabebe E, Anumba DOC. The Vaginal Microenvironment: The Physiologic Role of Lactobacilli. Frontiers in Medicine. 2018;5:181.

64. Bowman K, Rose J. Estradiol stimulates glycogen synthesis whereas progesterone promotes glycogen catabolism in the uterus of the American mink (Neovison vison). Animal Science Journal. 2017;88:45–54.

65. Nephew KP, Long X, Osborne E, Burke KA, Ahluwalia A, Bigsby RM. Effect of estradiol on estrogen receptor expression in rat uterine cell types. Biology of reproduction. 2000;62:168–77.

66. Fukuda M, Fukuda K, Ranoux C. Unexpected low oxygen tension of intravaginal culture. Human reproduction (Oxford, England). 1996;11:1293–5.

67. Edwards V, McComb E, Guttman H, Humphrys M, Forney L, Bavoil P, et al. P08.06 Lactic acid isomers differentially reduce <em>chlamydia trachomatis</em> infection in a ph dependent manner. Sexually Transmitted Infections. 2015;91 Suppl 2:A134 LP–A134.

68. Nardini P, Ñahui Palomino RA, Parolin C, Laghi L, Foschi C, Cevenini R, et al. Lactobacillus crispatus inhibits the infectivity of Chlamydia trachomatis elementary bodies, in vitro study. Scientific reports. 2016;6:29024.

69. Hearps AC, Tyssen D, Srbinovski D, Bayigga L, Diaz DJD, Aldunate M, et al. Vaginal lactic acid elicits an anti-inflammatory response from human cervicovaginal epithelial cells and inhibits production of pro-inflammatory mediators associated with HIV acquisition. Mucosal Immunology. 2017;10:1480–90.

70. Valore E v, Park CH, Igreti SL, Ganz T. Antimicrobial components of vaginal fluid. American Journal of Obstetrics and Gynecology. 2002;187:561–8.

71. Graver MA, Wade JJ. The role of acidification in the inhibition of Neisseria gonorrhoeae by vaginal lactobacilli during anaerobic growth. Annals of Clinical Microbiology and Antimicrobials. 2011;10:8.

72. Kumamoto Y, Iwasaki A. Unique features of antiviral immune system of the vaginal mucosa. Current opinion in immunology. 2012;24:411–6.

73. Fettweis JM, Serrano MG, Brooks JP, Edwards DJ, Girerd PH, Parikh HI, et al. The vaginal microbiome and preterm birth. Nature Medicine. 2019;25:1012–21.

74. Redelinghuys MJ, Geldenhuys J, Jung H, Kock MM. Bacterial Vaginosis: Current Diagnostic Avenues and Future Opportunities. Frontiers in cellular and infection microbiology. 2020;10:354.

75. Chee WJY, Chew SY, Than LTL. Vaginal microbiota and the potential of Lactobacillus derivatives in maintaining vaginal health. Microbial Cell Factories. 2020;19:203.

76. Cribby S, Taylor M, Reid G. Vaginal microbiota and the use of probiotics. Interdisciplinary perspectives on infectious diseases. 2008;2008:256490.

