## Supplemental Figures for "Vaginal microbiome-host interactions modeled in a human vagina-on-a-chip"

### SUPPLEMENTARY FIGURES

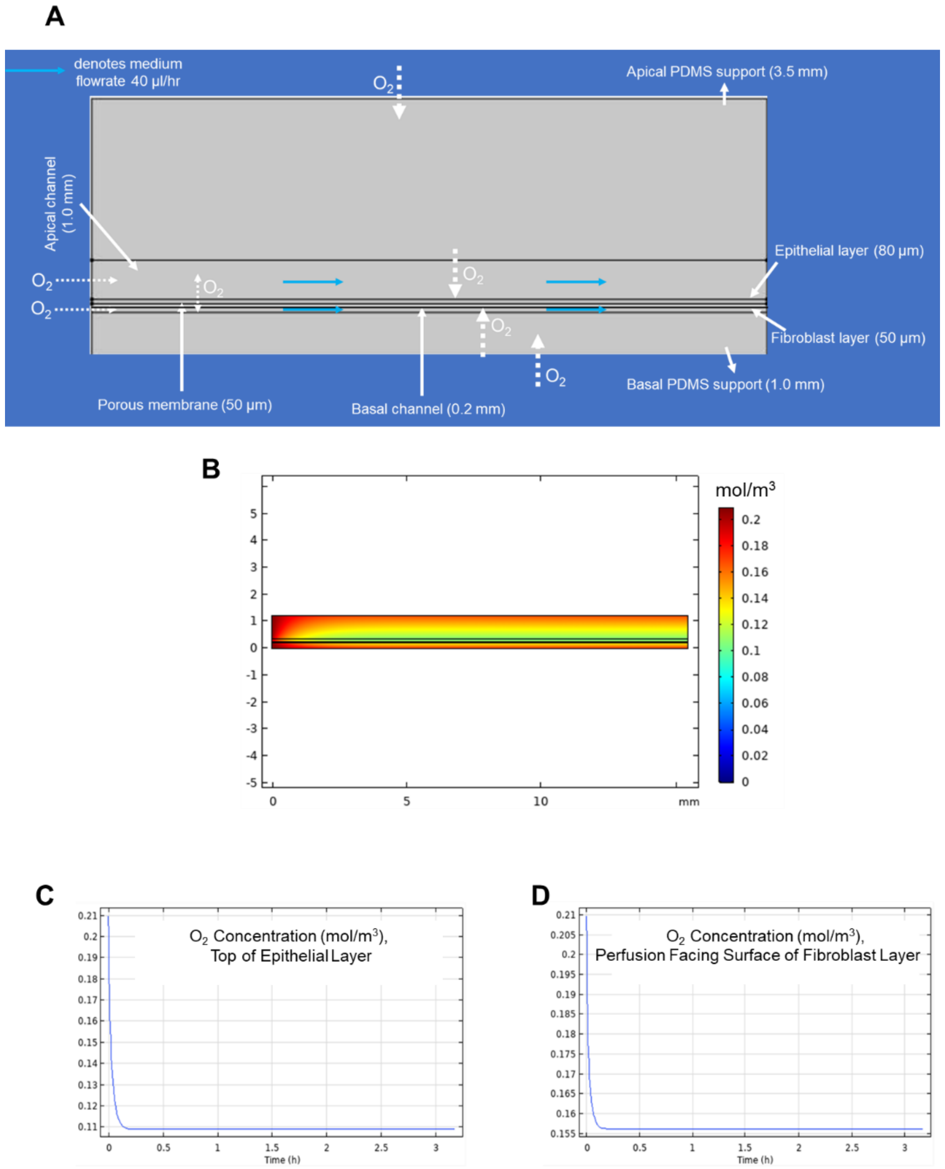


**Supplementary Fig. S1. Computational Model of Oxygen Gradient Generated by Vagina Chip.** COMSOL 2D model with geometry adapted from the commercial Organ Chip (from Emulate Inc.) used in these studies. The chip contains two parallel channels under continuous flow with thick epithelial and fibroblast cell layers cultured respectively on the top and bottom of a 50 µm thick porous membrane that separates the two channels. Dotted arrows show sources of oxygen inflow and consumption in the chip. **B)** Surface plot demonstrating $O_2$ distribution in the Vagina Chip. The lower graphs show results of $O_2$ concentration simulations over time in apical epithelial channel (**C**) and basal fibroblast channel (**D**).

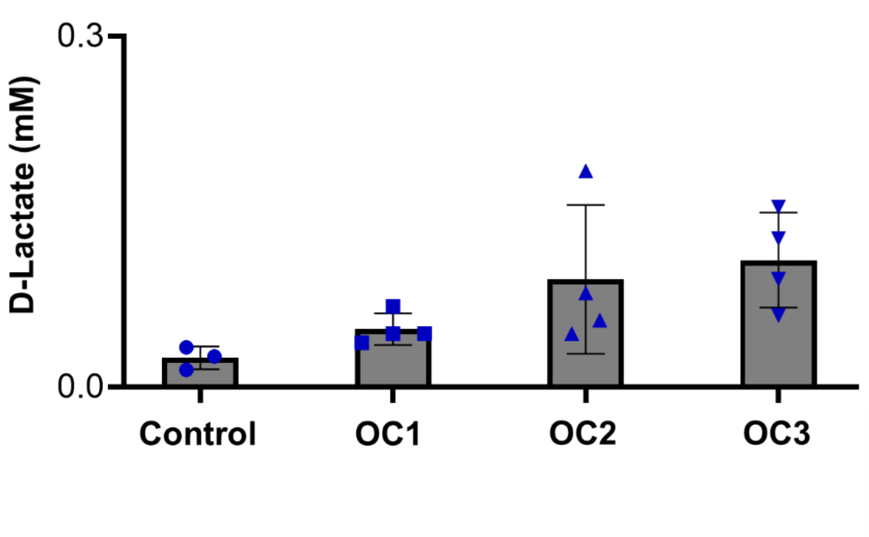


**Supplementary Fig. S2. D-lactate Production in Vagina Chips.** D-lactate concentrations measured in effluents from the apical epithelial channel of chips cultured in the absence (Control) or presence of the OC1, OC2, or OC3 *L. crispatus* consortia collected at 24 hours post inoculation are shown; each data point indicates one chip.
